# Fast turns are not sped up slow turns: How bluegill sunfish change their kinematics with turn speed

**DOI:** 10.1101/2025.01.24.634789

**Authors:** Andrew Clark, Eric Tytell

## Abstract

Fish turn extremely often, but this behavior is relatively understudied due to how challenging it can be to get fish to perform such an unsteady maneuver repeatedly. Specifically, little is known about whether fish control turns differently at different turning rates. Here we address the challenge of studying turning by developing a device that elicits turns repeatedly at specific speeds. Using this device, we compare the swimming kinematics of bluegill sunfish (*Lepomis macrochirus*) during fast and slow turns through 180 degrees. We find that the fish behave differently when turning quickly than when turning slowly: a fast turn is not a sped up slow turn, but is kinematically distinct. In particular, during fast turns, bluegill bend their bodies to minimize moment of inertia before they maximize torque, while in slow turns they maximize torque first, before they bend their bodies. In fast turns, they also beat their pectoral fins at a higher frequency, but take fewer pectoral fin strokes. Differences between the two speed turns may be due to initial momentum and how momentum is conserved throughout the turn, because fast turns have higher initial linear momentum that they can convert into angular momentum to turn around.

**Summary statement:** Bluegill change how they handle torque and moment of inertia when turning rapidly than when turning slowly. Tradeoffs between linear and angular momentum may play a key role.

## INTRODUCTION

Steady swimming in fishes has been studied extensively, but for many fish species, unsteady swimming comprises a large part of everyday locomotion. For example, trout spend most of their time accelerating, decelerating, and turning rather than swimming steadily (Webb, 1991). Similarly, both zebrafish and tuna, despite their large difference in size, swim unsteadily, turning regularly during normal swimming bouts (Fuiman and Webb, 1988; Howe and Astley, 2020; Thandiackal and Lauder, 2020; and Downs et al., 2023). Unsteady swimming becomes particularly important during predator-prey interactions, where both predator and prey accelerate, decelerate, and turn during encounters (Higham, 2007; see Webb, 1983; Webb, 1984; Webb, 1986; and Hitchcock et al., 2015 for examples). Fish also swim unsteadily during territory defense (e.g., Feldmeth, 1983). Finally, sharks swim unsteadily in social situations such that individuals lower in the hierarchy maneuver to keep out of the way of their superiors (Weihs et al., 1981) and will regularly turn with dramatic body bends (Porter et al., 2009).

In addition to being ubiquitous, understanding unsteady swimming is also important to understand the energy budget for fishes as they swim. On one side, muscles in some fish species fatigue less over time when they are used intermittently than when they are used continuously Coughlin and Dutterer (2024). In contrast, modeling of swimming states in rummy nose tetras suggests that burst and coast swimming uses more energy than continuous swimming (Ashraf et al., 2020) and *in vivo* studies in salmon show that they incur metabolic costs 40 times greater during burst swimming than during sustained swimming (Puckett and Dill, 1984), further highlighting the need to better understand the mechanics underlying unsteady swimming.

One unsteady behavior that has been studied extensively is the C start (Domenici and Hale, 2019). Importantly, the C start is often a strong response, and is straightforward to elicit with strong-enough stimuli. C starts consist of three stages: the preparatory, propulsive, and variable stage (Weihs, 1973). The first stage involves a rapid whole body bend into a “C” shape while the second stage is characterized by the return flip of the tail and subsequent acceleration Domenici and Blake (1997). C starts have high angular velocities and occur in very short time periods, often occurring in less than one fifth of a second (Domenici and Blake, 1997).

While C starts are important for survival, routine turning is a much more common behavior – but has been studied much less. From the current body of literature, we know that some fish initiate turning with an anterior bend while “planting” their tail in the water to use it is a fulcrum during the maneuver (Gray, 1933), an idea further supported by the finding that the anterior and posterior regions of the body generate large amounts of positive work at the initiation of turning in zebrafish (Thandiackal and Lauder, 2020). We also know that tuna can turn in one of three ways: with a gliding turn where they use their caudal fin as a rudder, with a powered turn where they keep beating their caudal fin symmetrically, and with a ratchet turn where they asymmetrically beat their caudal fin to one side of their body (Downs et al., 2023). Such asymmetry has also been reported in turning coelacanths (Fricke and Hissmann, 1992). Additionally, body caudal fin swimmers turn around when exposed to an oscillatory stimulus while median-paired fin swimmers do not (Marcoux and Korsmeyer, 2019). Finally, there have been some quantification of various physical quantities such as centripetal acceleration during turning(Howe and Astley, 2020).

Of particular note, Dabiri et al (2020) identified a mechanism that they proposed underlies routine turning across many species. They quantified physical quantities such as torque and moment of inertia as fish turned, concluding that timing is conserved across species so that animals keep their bodies long to maximize torque before bending their bodies to minimize moment of inertia and accelerate through the turn.

In this study, we examine the mechanism proposed by Dabiri et al (2020) to test whether it holds for turns at different speeds. Inspired by the system built by Marcoux and Korsmeyer (2019), we developed a system that allows us to repeatably elicit 180 degree turning behavior at controlled speeds. The system consists of a flow-through chamber attached to a movable platform that is mounted on top of a flow tank. Fish placed in the chamber reliably turn as the chamber changes direction. Using this system, we worked with bluegill sunfish (*Lepomis macrochirus*) to answer the question: are fast turns simply sped up slow turns, or are there distinct kinematic patterns that differentiate them?

When turning, fish face a tradeoff: they need to produce torque to propel themselves through the turn, but they can also reduce their moment of inertia to speed up the turn. In other words, by keeping their body straight, they generate larger amounts of torque using their tail because their lever arm is long, but they also have a larger rotational moment of inertia (Dabiri et al., 2020). Alternatively, by bending their body, they reduce their moment of inertia, but also reduce the lever arm for generating torque.

Dabiri et al. (2020) observed one solution for this tradeoff. They found that both zebrafish and jellyfish initially keep their bodies straight to produce large amounts of torque before bending their bodies to reduce moment of inertia, thus increasing their angular acceleration through the turn. Here, we explore whether these conclusions might change depending on the speed of the turn. In other words, do fish handle these tradeoffs differently when they need to turn quickly rather than when they have time to slowly turn around? We hypothesized that the tradeoffs would be handled differently: that a fast turn is not simply a sped up slow turn.

We ground this hypothesis in the equation for angular acceleration (*α*)

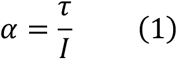

and the moment of inertia *I* for a body consisting of many segments is

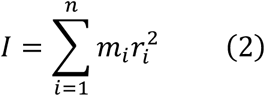

with *i* as the segment number, *m_i_* as the mass of the segment at a point, and *r_i_*^2^ as the distance from that mass to the center of rotation. Torque *τ* is

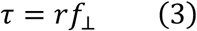

where *r* is the distance from the center of rotation to the propulsor and *f*_⊥_ is the force the propulsor produces that is perpendicular to *r*.

Based on (Equation 1), to have a higher angular acceleration, the fish can either increase torque or decrease moment of inertia. Both the torque and moment of inertia depend on *r*, but while torque scales with *r*, moment of inertia scales with *r*^2^ (Equation 2 and Equation 3).

Therefore, by bending its body, the fish linearly decreases torque, but quadratically decreases moment of inertia. Since angular acceleration needs to be higher during faster turns, we thus hypothesized that fish would prioritize body bending in fast turns to reduce moment of inertia, but would prioritize high torque by keeping the body straight at the beginning during slow turns, similar to what Dabiri (2020) observed.

Additionally, we quantified linear and angular momentum of the body during the turn to examine how fish exchange linear and angular momentum, and whether this exchange might differ depending on speed. Finally, we examined pectoral fin movements during the turns. Bluegill use their pectoral fins regularly for maneuvering and stability Drucker and Lauder (2001). We expected that turns at different speeds might have different stability or control requirements, and this would be reflected in differences in pectoral fin use.

## METHODS

### Setup

Inspired by the system used by Marcoux and Korsmeyer (2019) we developed an oscillatory device to repeatably induce unsteady behaviors (Figure 1 A,B). We mounted a Teknic ClearPath-SDSK servo motor (part number CPM-SDSK-2310S-EQN) to an 80/20 chassis with an attached acrylic flow-through chamber that is 20.3cm wide by 30.5cm long and 7.6cm tall. When on top of the flow tank, the water level sits about 20.5cm above the bottom of the flow through chamber. This device drove forwards and backwards at programmable rates controlled by an Arduino Uno. The flow-through chamber had metal grates on either end to keep the fish inside and translucent plastic sheets taped on either side with vertical lines drawn on them to provide a visual stimulus for the fish. The wheels of the car ran in a metal track. Since the fish is contained by the grates of the car, when the car reverses direction, the fish must turn at roughly the same rate.

**Figure 1:**
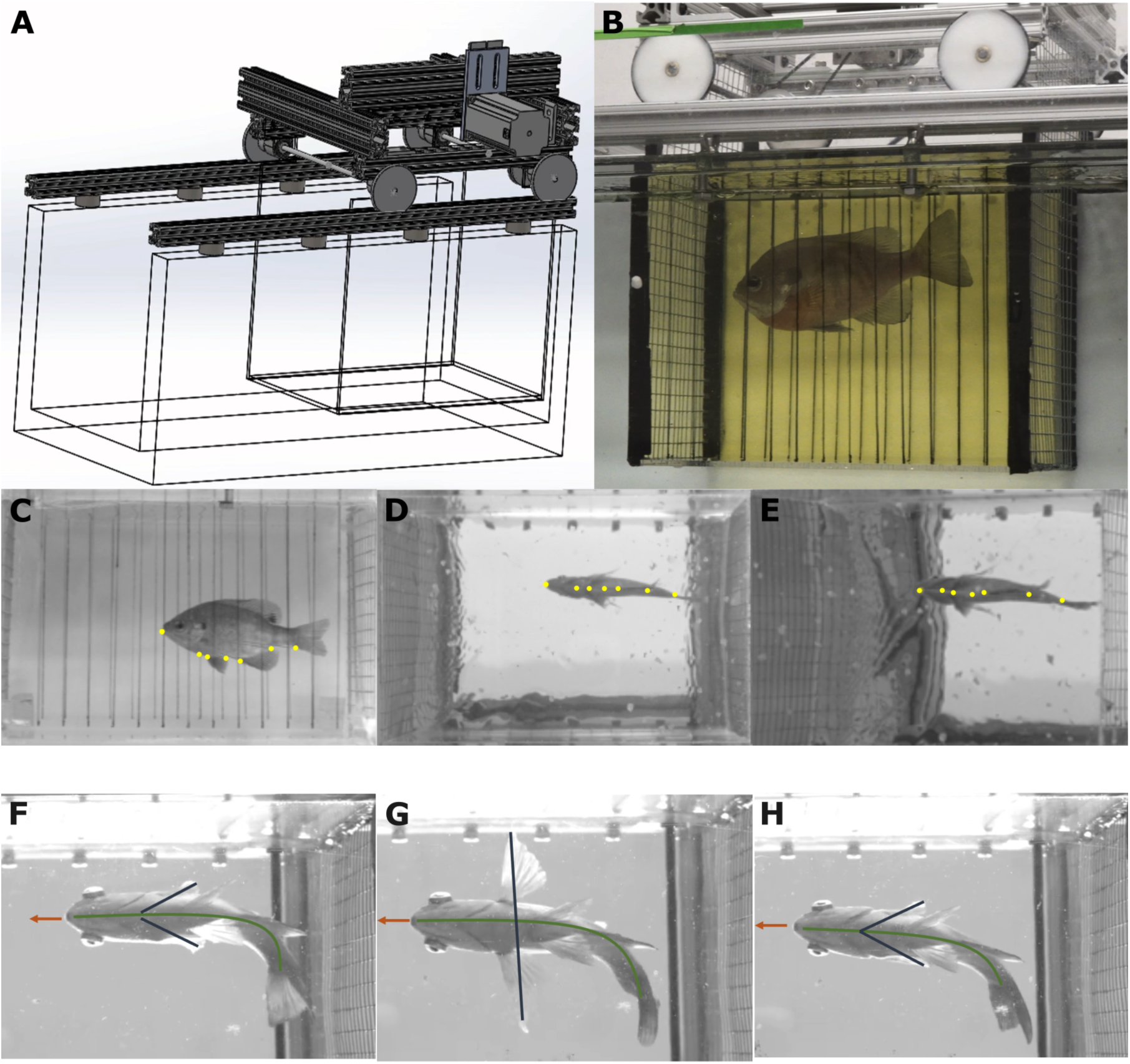
Our programmable car elicits repeatable turns. (A) A 3D solidworks model of the system (available in supplementary material). (B) A bluegill sunfish swimming in the chamber. (C, D, E) We filmed using an orthogonal lateral camera (C), an orthogonal ventral camera (D), and an off axis ventral camera (E). The yellow points represent digitized points along the body. Light gray lines show the skeleton used for training (the snout to all other points to form a shallow tree). (F, G, H) Three consecutive turns show similar kinematics from the ventral view.

These turns were generally repeatable (see Figure 1 C-E for an example) and their consistency allowed us to collect multiple trials with similar performance. For all trials, we kept the water still while the car moved along the track and ran the car at either a low speed (around 1 body lengths *L* per second at peak speed) or a high speed (either 2.5 or 3.5 *L*/sec at peak speed) to elicit slow and fast turns respectively. When the fish refused to remain centered in the flow through chamber, it was gently encouraged to move with a wooden dowel. We defined the start of each turn as the first observable movement of the snout in the direction of the turn, and the end as when the fish’s snout was oriented in the opposite direction (an approximately 180 degree turn).

### Filming and digitizing

To film the kinematics, we set up two Phantom Miro M340 high-speed cameras and one Phantom Miro M120 high-speed camera. We positioned two cameras orthogonal to each other: one directly below the working section of the tank and one directly lateral to the working section of the tank, with one positioned ventrally and at an angle to the rear to the other cameras. We filmed all videos at 60 frames per second. We calibrated videos by using a ChArUco board (Garrido-Jurado et al., 2014) and the aniposelib package from the Anipose toolkit (Karashchuk et al., 2021).

We used SLEAP (Pereira et al., 2022) to digitize seven points along the midline on the bottom of the fish: the snout, a point in line with the posterior portion of the gills, a point between the pelvic fin insertions, a point halfway between the pelvic fin insertions and the anal fin insertion, the anal fin insertion, a point halfway between the anal fin insertion and the peduncle, and the peduncle (Figure 1 C-E). We wrote custom code to use a ChArUco board to calibrate the filming space and triangulated all digitized points from SLEAP into 3D coordinates in centimeters. We then rotated all points such that they conformed to axes defined by the flow-through section of the apparatus, where the *x* axis is forward, the *y* axis is to the fish’s left, and the *z* axis is up. Finally, we used the projections of the 3D points into the *x*, *y* for all analyses, since those were the primary axes in which the turns took place. This allowed us to look at the turns in the important axes while correcting for parallax introduced if the fish was at different vertical locations.

We smoothed the *x* and *y* locations of all digitized points individually using a 9th order low pass Butterworth filter with a cut off frequency of 8 Hz for fast turns and 4 Hz for slow turns. We then used a cubic smoothing spline to spline the points along the body, interpolating locations that are evenly spaced along the body.

### Snout and tail angle and angular velocity

We estimated snout angle by first drawing a line from the center of the pelvic fin insertions to the snout, and tail angle similarly with peduncle and the point halfway between the base of the anal fin and the peduncle. The angle between this vector and the horizontal *x* axis (*θ*) is then *θ* = atan2(*y*_1_ – *y*_2_, *x*_1_ – *x*_2_), where *y*_2_ and *x*_2_are the y and x positions of the snout or peduncle and *y*_2_ and *x*_2_ are the y and x positions of the pelvic point or anal fin point, respectively.

We then smoothed the angles using a low pass Butterworth filter with a threshold of 4 Hz before numerically differentiating the angle with respect to time by estimating the central difference between points (Smith, 1985). Positive angular velocity values denote movement in the direction of the turn while negative angular velocity values denote movement in the direction opposite the turn.

### Moment of inertia

To estimate the mass of each segment along the body, we first estimated their volumes using FIJI (Schindelin et al., 2012) to measure height, width, and length of each segment of the fish. We then estimated volume by approximating each segment as a truncated elliptical cone and mass by assuming the fish’s density is nearly constant. Here, *w* and ℎ are width and height of the fish’s body segments, with *w* as the blue lines in Figure 2 b and ℎ as the blue lines in Figure 2 a. We drew each segment line halfway between two adjacent points down the length of the body, with the exception of the snout and peduncle point where we drew one segment line at the point. For each point along the body, we used the width and height of the body segment line directly anterior to the point (*w_i_* and ℎ*_i_*) and the width and height of the body segment line directly posterior to the point (*w_i_*_+1_ and ℎ*_i_*_+1_).

**Figure 2:**
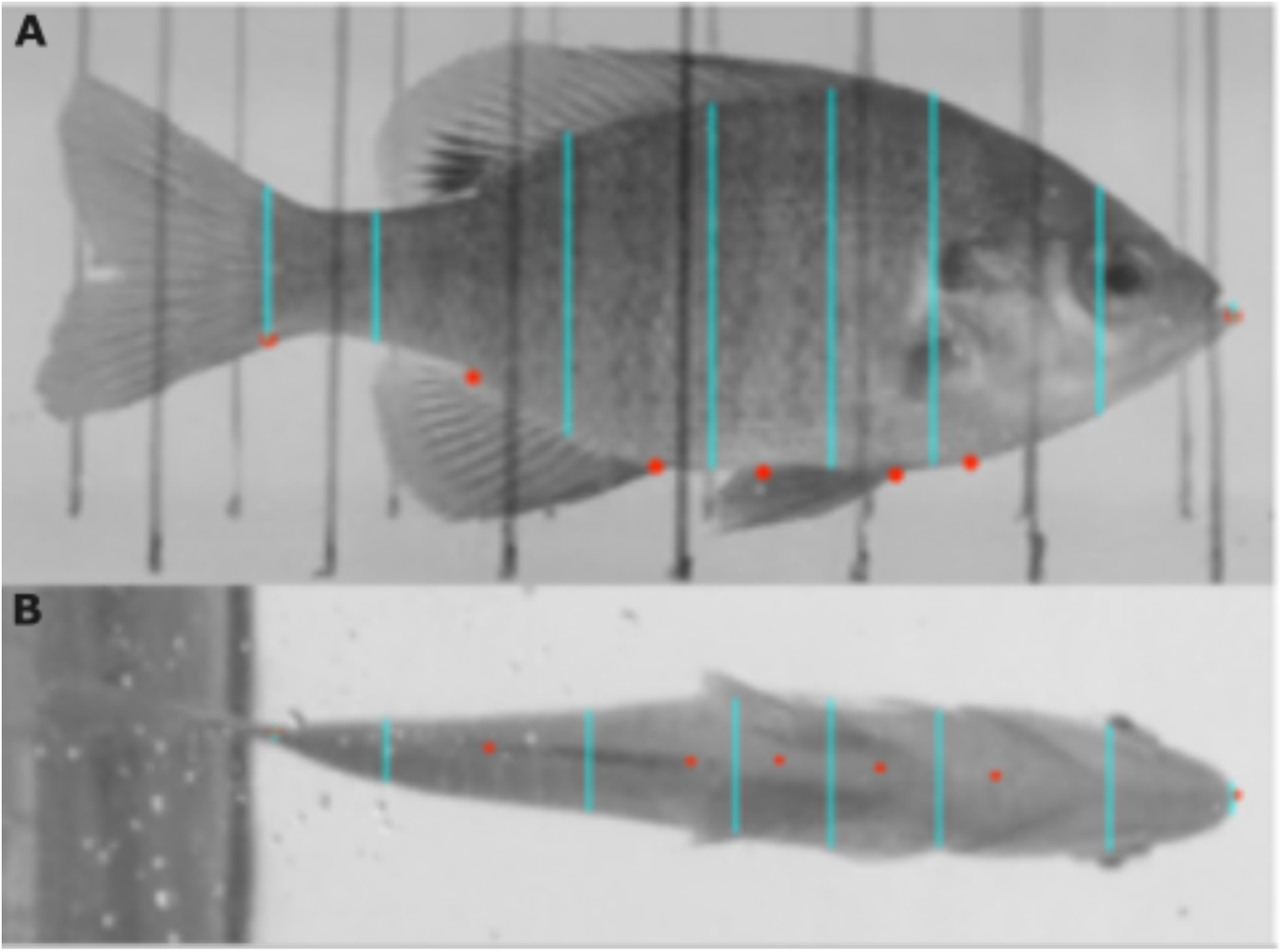
Mass was estimated using images of each bluegill. (A, B) Points were added to their respective locations along the length of the body (red dots). Lines were then drawn from vertical edge to vertical edge of the fish in the lateral view (A) and lateral edge to lateral edge of the fish in the ventral view (B) (light blue lines). FIJI was then used to measure each of the lines and the distance between them in the ventral view. The known length of the fish was used to set the scale.

The volume *V_i_* of each segment is thus

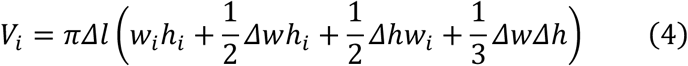

where *Δl* is the distance between segments, *Δw* = *w_i_*_+1_− *w_i_* and *Δ*ℎ = ℎ*_i_*_+1_ − ℎ*_i_*, and the mass of each segment is then *m_i_* = *ρV_i_*. Finally, we approximated each segment’s mass as a point mass located at the digitized points splined along the body such that the distances between them were consistent with the known distances along the body of the fish.

The moment of inertia (#eq-moi) was then estimated, assuming the center of rotation was the whole body mass centroid (*X_M_*, *Y_M_*)

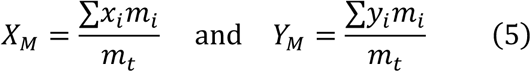

where *m_i_* is the mass of each segment. We normalized moment of inertia by dividing by the moment of inertia for each individual when its body was straight.

### Torque

We estimated the whole body torque of the fish based on of the sum of the torques of the segments. In a non-rigid body, torque is the time derivative of moment of inertia of each body segment (*I_i_*) multiplied by its corresponding angular velocity (*ω_i_*):

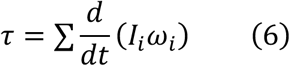

The moment of inertia *I_i_* for each segment is equal to the segment’s mass multiplied by the square of its distance to the center of rotation (*r_i_*). Further applying the chain rules gives us the whole body torque of the fish:

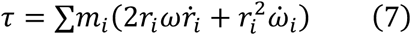

### Pectoral fins

We manually digitized the start of all pectoral fin strokes that occurred during the turn. Working with only ventral orthogonal videos, we identified the frames in which each pectoral fin was maximally extended away from the body, immediately before the fish pulled it back towards the body to take a stroke. We digitized the tips of each pectoral fin when it was extended in a frame and the bases of both pectoral fins in all frames in which either fin was extended. In many instances, bluegill seem to be taking “backing” strokes in which they pushed their pectoral fins far towards their snout to produce a backwards force on that side. For our analyses, we considered every pectoral fin stroke greater than 90 degrees (maximally extended) to be a backing stroke.

### Momentum

We estimated the linear momentum *p*_*_ of the fish during turning as

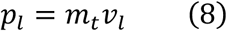

where linear velocity *v*_*_ is the central difference of the position of the whole body mass centroid differentiated with respect to time and *m*_)_ is the total mass of each fish. Because maximum velocity tends to be proportional to body length *L*, and body mass is proportional to *L*^3^, momentum should be proportional to *L*^4^. To control for individual size differences, we therefore divided momentum by *L*^4^. We estimated angular momentum *p_θ_* as

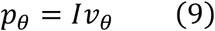

where *I* is the moment of inertia we estimated using Equation 2 and *v_θ_* is the angular velocity of the fish’s snout as estimated above.

For both linear and angular momentum, we estimated the pre-turn momentum, defined as the mean of the momentum in the two frames leading up to the turn, the mean momentum during the first half of the turn and, and mean momentum during the second half of the turn. We divided the turns into halves according to total time of each turn.

### Statistics

For all analysis and visualization, we used R version 4.3.3 with Rstudio version 2023.12.1.402 and the following packages: tidyverse version 2.0.0 (Wickham et al., 2019), lme4 version 1.1.35.1 (Bates et al., 2015), car version 3.1.2 (Fox and Weisberg, 2019), ggspatial version 1.1.9 (Dunnington et al., 2023), ggdist version 3.3.2 Kay and Wiernik (2024), and beeswarm version 0.4.0 (Eklund and Trimble, 2021).

When we use pairwise comparisons to look at the effect of the “slow” and “fast” speed groups, or to look at the significance between changes in momentum and turn portion, we use the Anova function from the car package and the lmer function from the lme4 package to run an ANOVA on a linear mixed-effects model with individual as a random effect. When we present linear and exponential fits, we similarly use the lmer function and fit the 1/x curve for Figure 9 C using ggplot2’s geom_smooth function.

All means are given plus or minus standard deviation, unless otherwise noted.

## RESULTS

We collected 5 bluegill sunfish (*Lepomis macrochirus*) (of lengths 12.5cm, 13cm, 14.5cm, 19cm, 19cm) from either White pond or Walden pond in Concord, MA. Each individual contributed at least 15 slow and 15 fast turns to the dataset.

### Angular velocity is higher when the car reverses faster

During fast car reversals (which we will call “fast turns”), fish turned 180 degrees in less time than they did during slow car reversals (“slow turns”), resulting in higher angular velocities (Figure 3). For each turn, we took the mean and maximum angular velocity over the duration of the turn. Then, we examined the means of these values across all turns. Both mean of the mean and the mean maximum snout angular velocity were significantly higher in fast turns (mean = 181 ± 41 deg/s; max = 428 ± 84 deg/s) relative to slow turns (mean = 96 ± 35 deg/s, max 303 ± 89 deg/s) (for both mean and max, p < 0.001).

**Figure 3:**
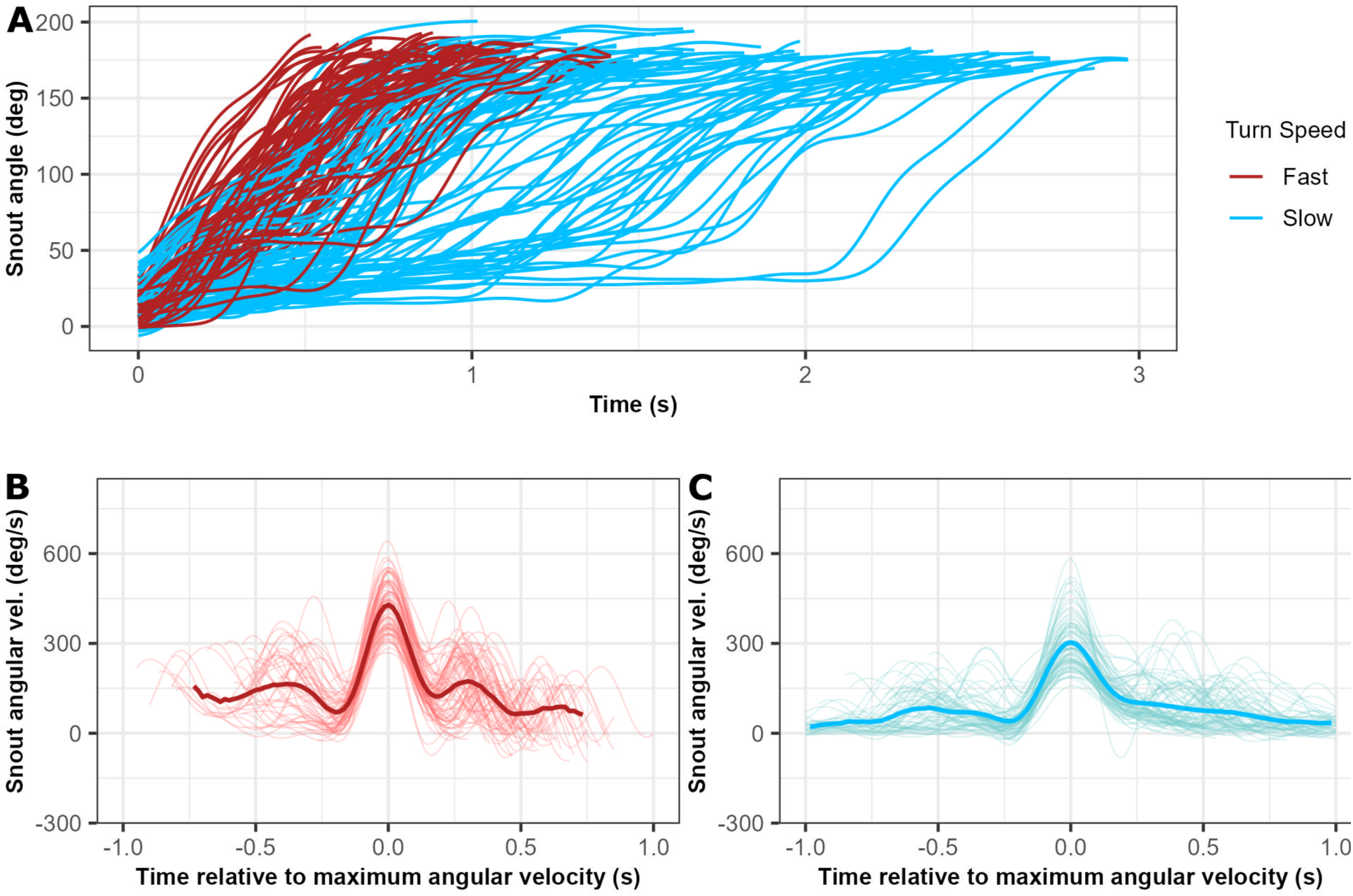
Angular velocities are higher when the car reverses faster. (A) Changes in snout angle for both slow turns in blue and fast turns in red using degrees over seconds. (B, C) Angular velocity with slow turns in blue (B) and fast turns in red (C)

### Changes in snout angular velocity are driven by anterior bending

We found that when the snout is moving at its maximum angular velocity, the tail is moving at a significantly lower angular velocity (p < 0,001) and is close to 0 (Figure 4 B). The reverse is also true: when the tail is moving at its maximum angular velocity, the snout is moving at a significantly lower speed (p < 0.001) that is close to 0 (Figure 4 C).

**Figure 4:**
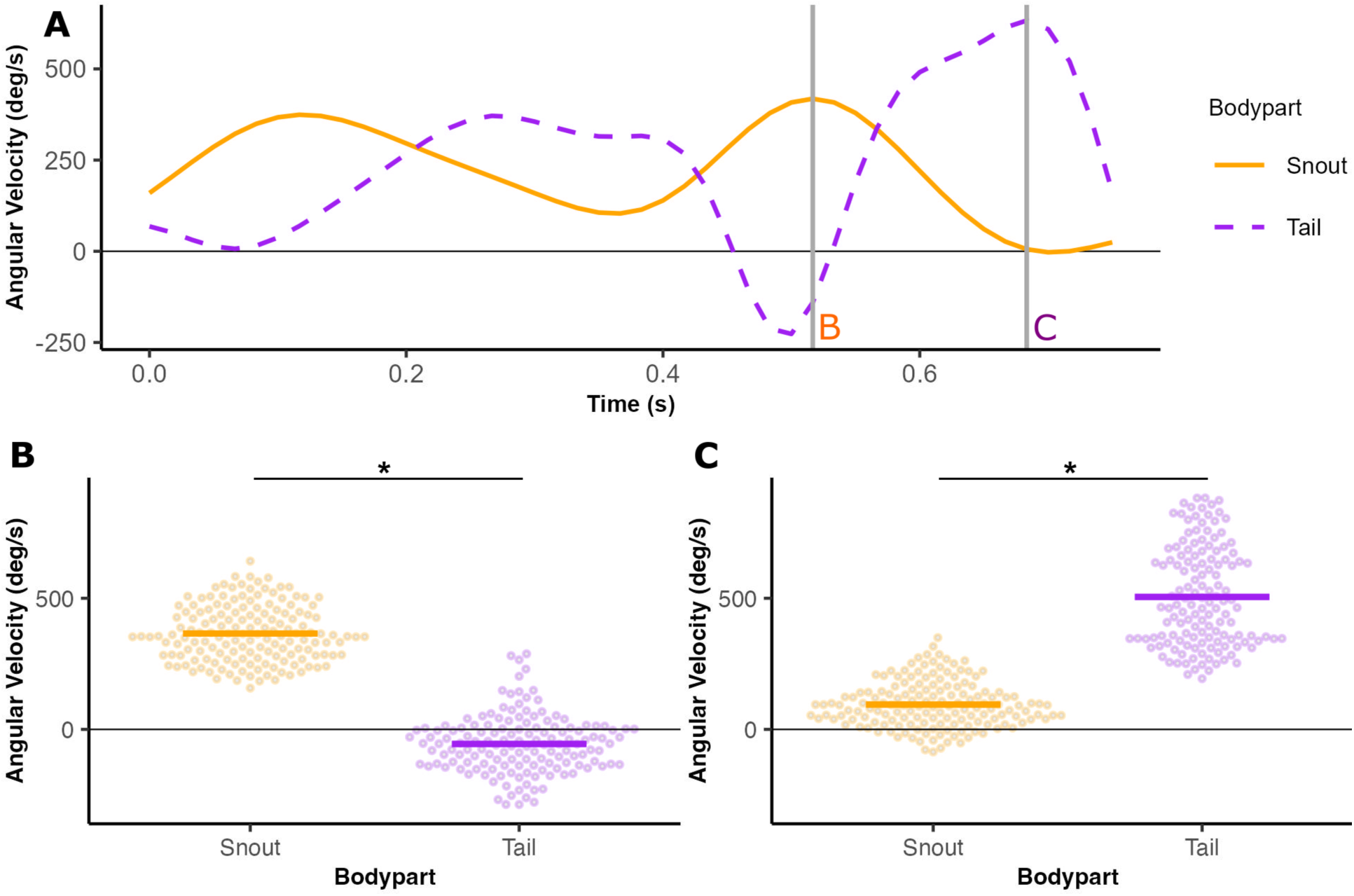
(A) Representative traces of one trial’s angular velocities over time for the snout (orange) and tail (purple). The gray lines represent the time of maximum angular velocity for the snout (orange “B”) and tail (purple “C”). (B) Snout and tail angular velocities for each trial at the point where snout angular velocity is maximized (first gray line in A). Each point denotes the value for one trial. (C) Snout and tail angular velocities at the point where tail angular velocity is maximized (second gray line in A).

Additionally, for all turns, fish asymmetrically beat their caudal fin and also simultaneously beat the median fins to that same side as the caudal fin. In about 1/3 of fast turns, fish bend and hold their median fins to the side of the turn before beginning to beat their tails.

### The minimum moment of inertia is similar for slow and fast turns

Whether fish turned slowly or quickly, they bent their bodies by about the same amount, decreasing their moment of inertia to about the same value, despite different turning rates (Figure 5). The mean value of the minimum normalized moment of inertia across the duration of a slow turn was 0.9 ± 0.04 and for fast turns was 0.9 ± 0.03 (p = 0.9056).

**Figure 5:**
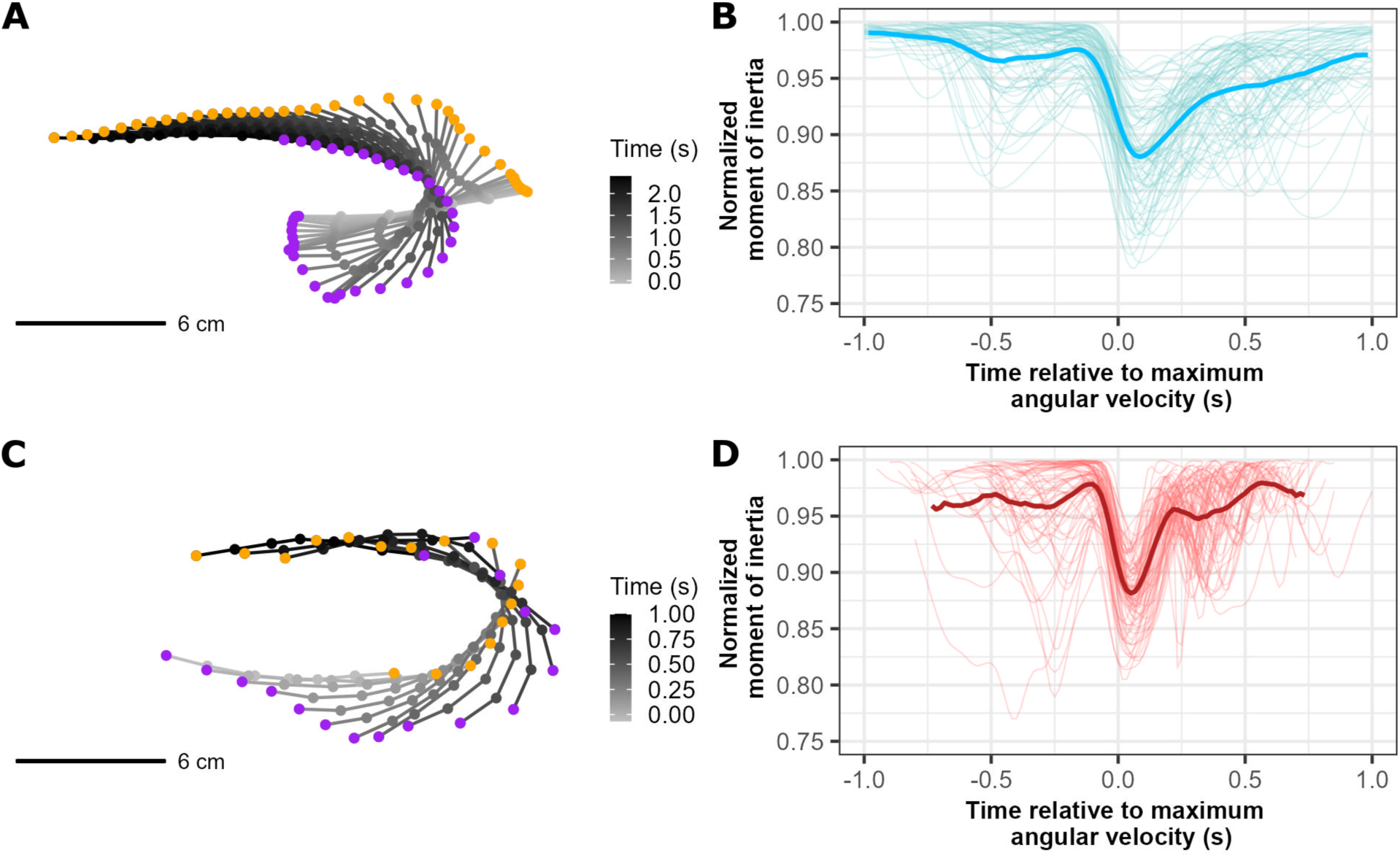
Fish bend their bodies to a similar degree in slow and fast turns. (A, C) Body traces of midlines for one individual for one slow (A) and one fast (C) turn. Points along the midlines denote digitized points, where the orange point is the snout and the purple point is the peduncle. Line color in (A) and (C) corresponds to the time course of the turn with darker gray representing the beginning and lighter gray representing the ending. Note the different time scale between fast and slow turns. (B, D) Normalized moment of inertia for slow (B) and fast (D) turns. Time is centered on the time of maximum angular velocity.

### Lower moments of inertia result in higher angular velocities regardless of turn speed

Overall, fish that bent their bodies more or held the bend longer over the course of the entire turn (resulting in a lower mean moment of inertia) turned with higher mean angular velocities (Figure 6). Fitting with a linear mixed effects regression with individual as a random effect yields a significant negative slope for both slow (p < 0.001) and fast (p < 0.001) turns. The slopes are the same for both types of turns (p = 0.4083). As with minimum moment of inertia, the mean moment of inertia over the entire turn did not differ significantly between slow and fast turns (0.95 ± 0.02 for both, p = 0.1136), but the mean angular velocity is higher in fast turns (Figure 3).

**Figure 6:**
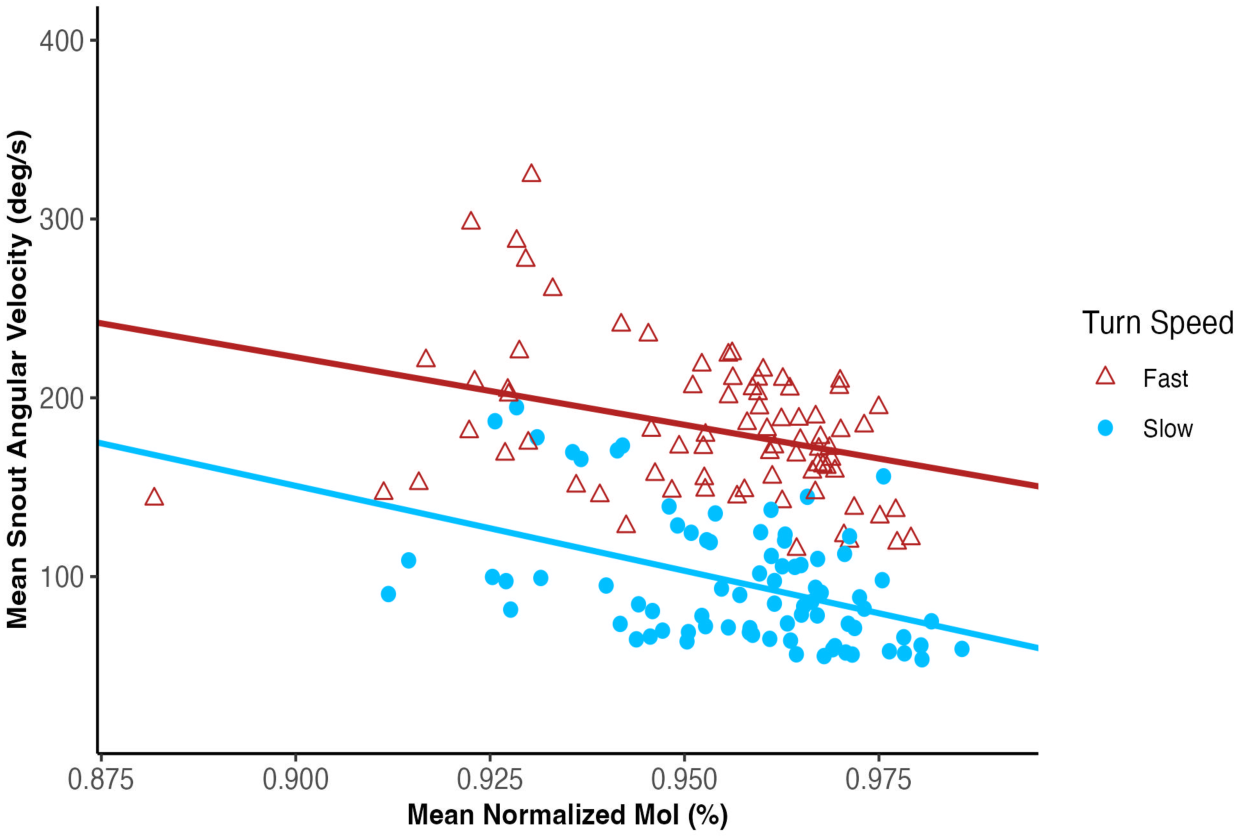
Bluegill that bend their bodies more generate higher angular velocities. Mean moment of inertia per trial, normalized by the maximum moment of inertia for that trial, plotted against mean snout angular velocity of that trial. Data is shown for both slow (in blue) and fast turns (in red). Linear regressions for slow (light blue) and fast (red) turns are shown as dashed lines.

For the vast majority of trials (130 of 139), fish reached their maximum angular velocity before they fully minimized their moment of inertia (points in the bottom right shaded region in Figure 7). Both slow and fast turns followed this pattern with few exceptions.

**Figure 7:**
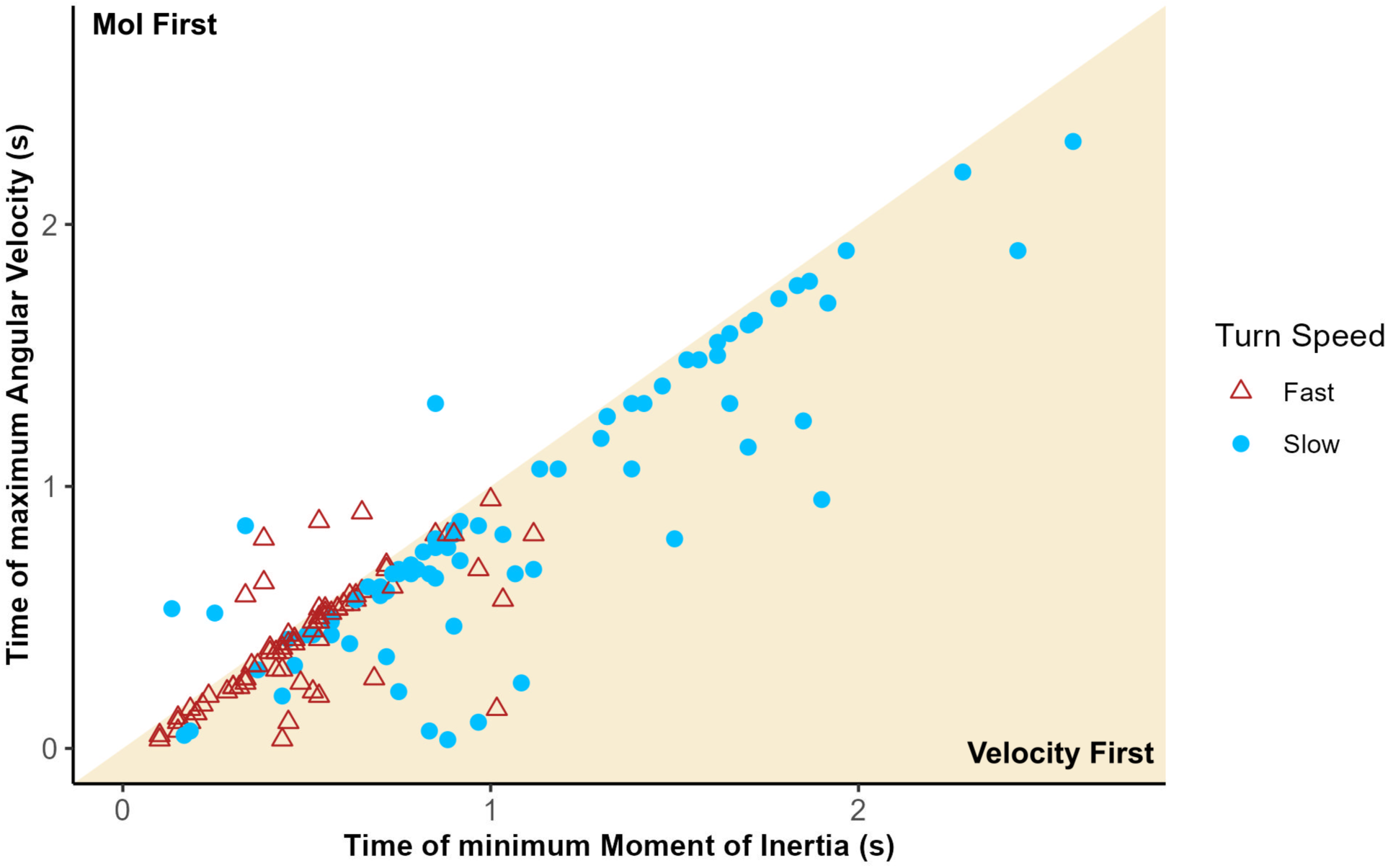
Bluegill consistently maximize angular velocity before minimizing moment of inertia. The time at which moment of inertia is minimized during a turn is plotted against the time at which angular velocity is maximized during that turn for both slow (blue) and fast (red) turns. The yellow shaded region is where the fish maximizes its angular velocity before minimizing its moment of inertia, while the unshaded region is where the fish minimizes its moment of inertia before maximizing its angular velocity.

### Fast turns trade off linear and angular momentum

Linear momentum differs between the two types of turns (Figure 8 A, C), while angular momentum shows a similar pattern across the turns. Pre-turning linear momentum is significantly higher in fast turns than it is in slow turns (p < 0.001). For fast turns, linear momentum then significantly *decreases* in both the first and second half of the turn (p < 0.001 in both cases). However, for slow turns, it does not change significantly in the first half of the turn (p = 0.12) and significantly *increases* in the second half of the turn compared to pre turning phase (p < 0.001). In other words, in fast turns, bluegill initially start with higher linear momentum, which decreases during the entire turn, corresponding to an increase in angular momentum (Figure 8 B). In slow turns, bluegill initially start with low linear momentum (Figure 8 A), increase angular momentum in the first half of the turn while maintaining linear momentum, then increase linear momentum in the second half of the turn as angular momentum decreases (Figure 8 a, B).

**Figure 8:**
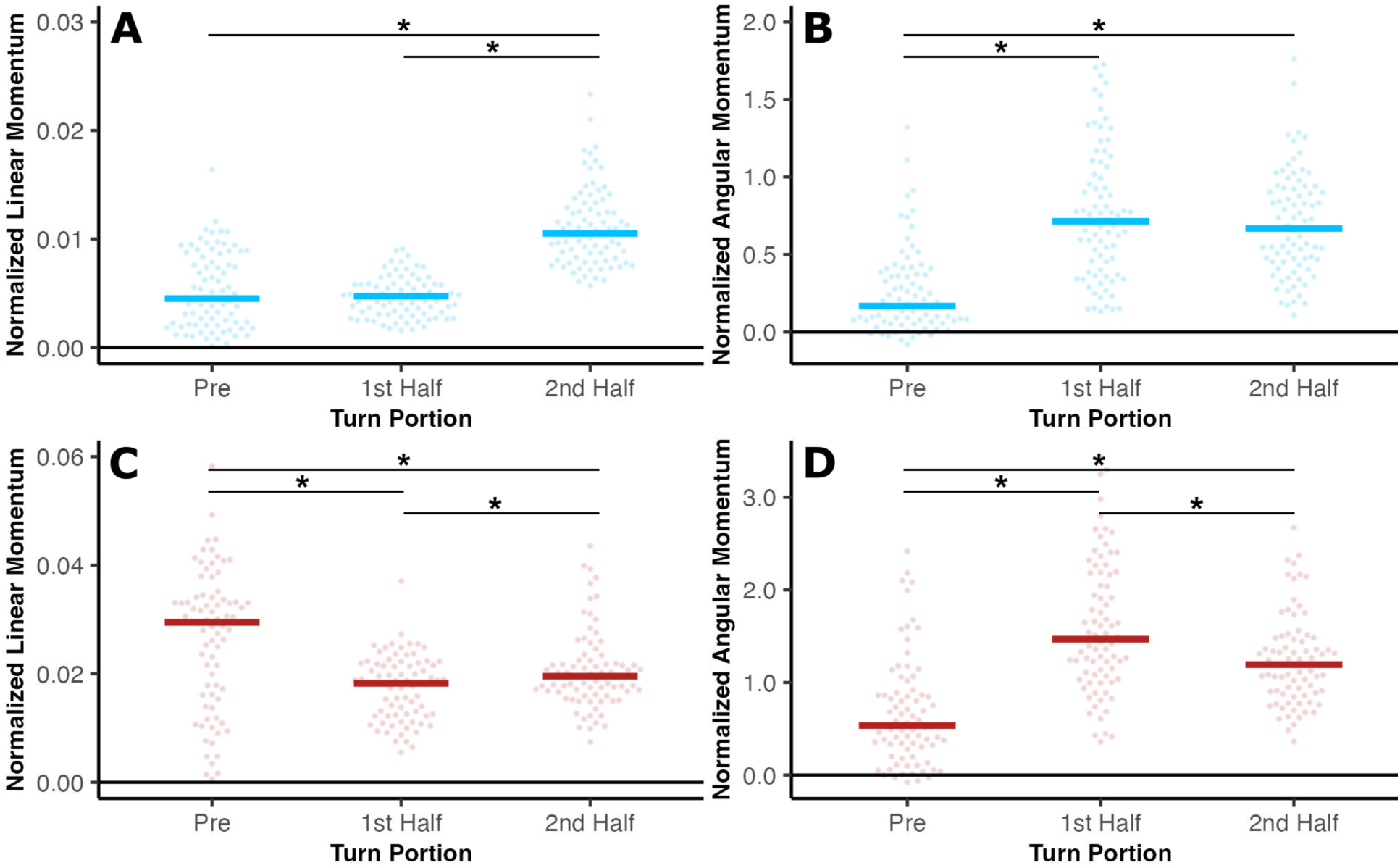
Bluegill convert linear to angular momentum in fast turns but not slow ones. (A,B) Linear (A) and angular (B) momentum for slow (blue) turns before the turn, during the first half of the turn, and during the second half of the turn. (C,D) Linear (C) and angular (D) momentum for fast (red) turns before the turn, during the first half of the turn, and during the second half of the turn.

### Bluegill use their pectoral fins differently during fast turns

Bluegill take more strokes with their pectoral fins in slower turns (based on a significant positive slope on the regression: p < 0.001) (Figure 9 A), but beat their pectoral fins at a higher mean frequency in fast turns (based on a significant inverse relationship on the regression: p < 0.001) (Figure 9 B). During slow turns, fish beat both their inside and outside pectoral fins at a similar amplitude (p > 0.3622), but during fast turns, fish beat their inside fin at a higher amplitude than their outside fin (p < 0.001) (Figure 9 C). The inside fin amplitudes do not differ between slow and fast turns (p = 0.6432) (Figure 9 C). Finally, bluegill take fewer backing strokes during fast turns (p < 0.001) (Figure 9 D). All individuals used their pectoral fins in all turns.

**Figure 9:**
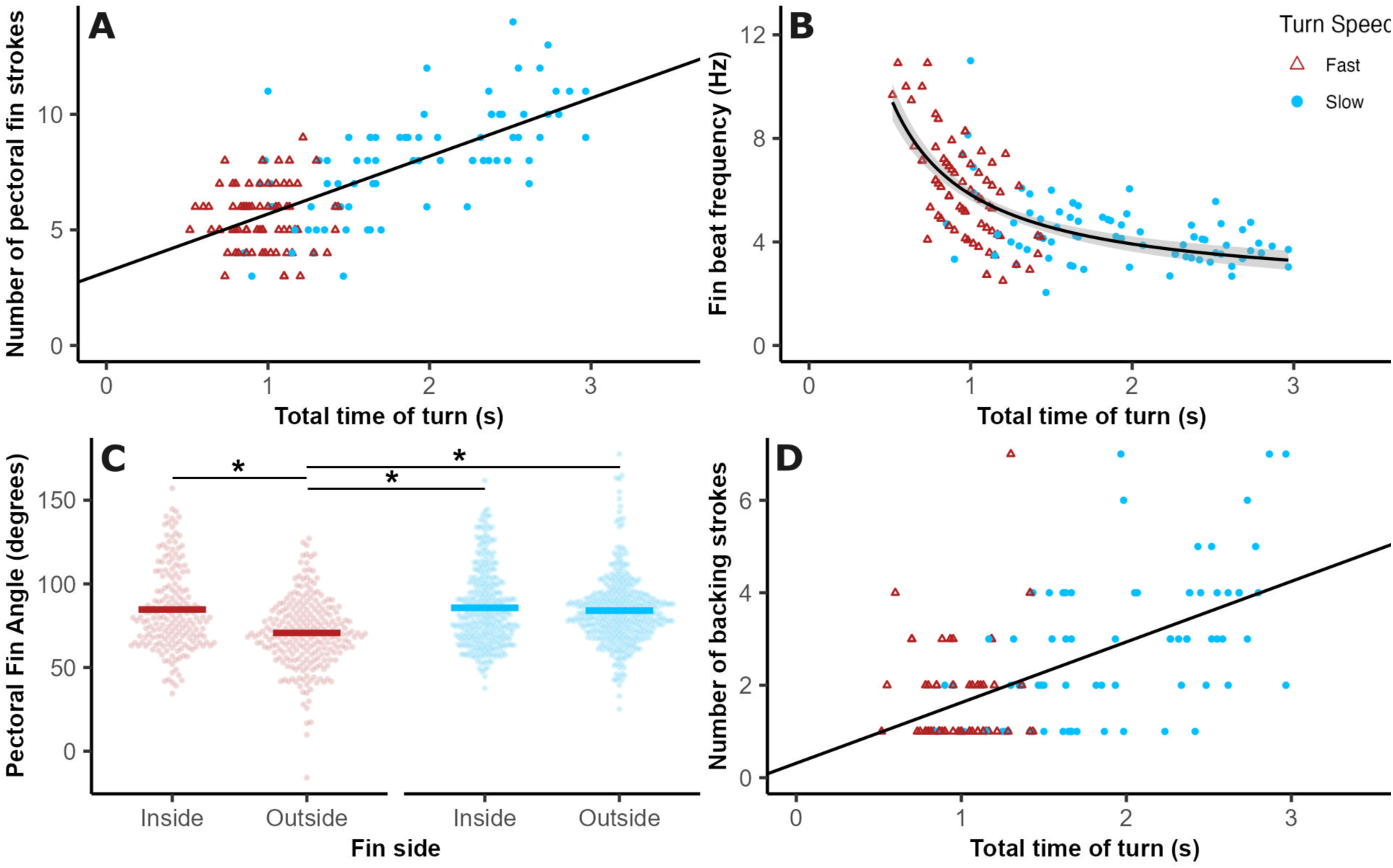
Bluegill use their pectoral fins differently at different speeds. Pectoral fin data for slow (blue circles) and fast (red triangles) turns. (A) Total number of pectoral fin beats increases for longer turns. The black line is the linear fit. (B) Fin beat frequency decreases for longer turns with a 1/x model shown by the black line. (C) Fin amplitude for the inside and outside fins in slow and fast turns. (D) Number of backing strokes increases for longer turns. The black line shows the linear fit.

### Timing of maximum torque and minimum moment of inertia differs based on turn speed

The timing of the estimated maximum torque relative to moment of inertia was distinctly different for slow and fast turns. For 51 of 79 slow turns fish maximized torque before minimizing moment of inertia. In contrast, for 54 of 78 fast turns, they minimized moment of inertia first. For slow turns, fish minimized their moment of inertia a mean of 0.06 ± 0.3s *after* maximizing torque, while for fast turns, fish minimized moment of inertia 0.2 ± 0.3s *before* maximizing torque (p < 0.001) (Figure 10).

**Figure 10:**
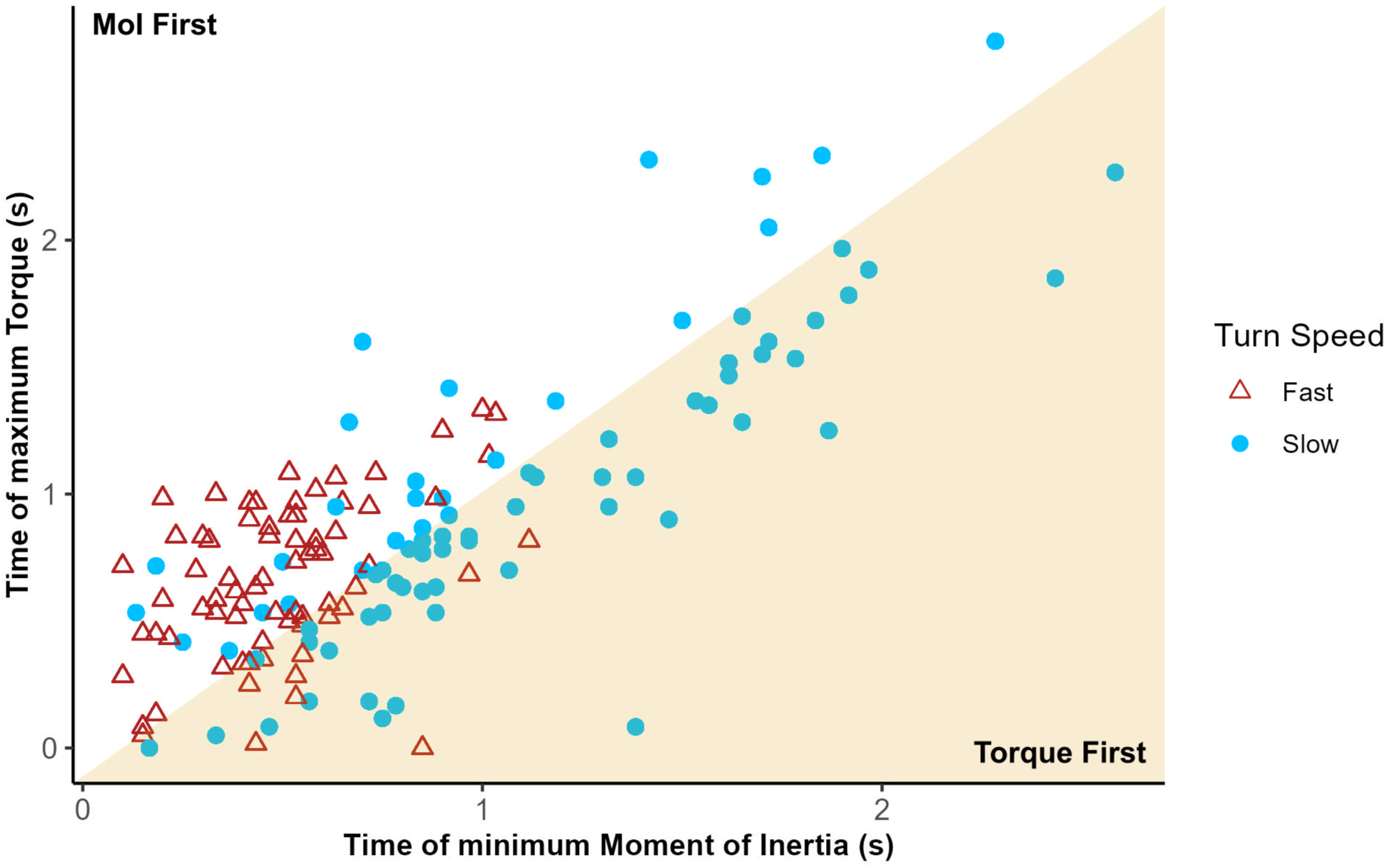
Slow and fast turns differ in their timing of maximum torque and minimum moment of inertia. The time at which moment of inertia is minimized during a turn is plotted against the time at which torque is maximized during that turn for both slow (blue) and fast (red) turns. The yellow shaded region is where the fish maximizes its torque before minimizing its moment of inertia, while the unshaded region is where the fish minimizes its moment of inertia before maximizing its torque.

## DISCUSSION

### Fast turns are not sped up slow turns

Fast turns are not sped up slow turns, but are kinematically distinct. During slow turns, bluegill behave similarly to jellyfish and zebrafish as studied by Dabiri et al (2020), reaching similar angular velocities and maximizing torque before minimizing moment of inertia. However during fast turns, bluegill do something different, reaching higher angular velocities and maximizing torque after minimizing moment of inertia. This change in timing coupled with the discrete differences in how linear momentum changes during turning suggests that fish change their kinematics based on some threshold of turn speed (Figure 10 and Figure 8).

Physically, it makes sense that bluegill change their timing of torque and moment of inertia. Due to how torque scales with *r* (Equation 3), while moment of inertia scales inversely with *r*^2^ (Equation 2), a reduction in *r* decreases moment of inertia more than torque. Therefore, bluegill prioritize minimizing moment of inertia early to maximize angular acceleration at the start. For slower turns, they can instead turn how Dabiri et al (2020) found and maximize torque at the beginning with a straighter body before bending to accelerate through the turn.

Our results contrast with those of Howe and Astley (2020) in that we find discrete differences based on speed while they do not. We believe this is because all of the turns they studied would fall into our category of “fast turns”. They did not control the heading change, and therefore most of the turns they studied were much smaller than those presented here. The largest was 120 deg (Howe and Astley, 2020), which is below the 180 degree turns we studied. However, even the turns that had a relatively small angular change had relatively high angular velocities, and would fall into our fast turn category. Their routine turns had maximum angular velocities of around 600 deg/s, which is greater than the maximum angular velocities in our fast turns, which averaged around 430 deg/s.

Our slow turns then represent a qualitatively different behavior than fast turns, and than the turns studied by Howe and Astley (2020). We believe they represent the type of behavior that fish might use to maneuver in a controlled manner around an obstacle or while grazing. For example, coelacanths make similar slow turns when maneuvering around an obstacle (Fricke and Hissmann, 1992).

### Fast turns are not C starts

Importantly, the turns we measured are not C starts. Compared to C starts, even our fast turns take much longer to complete. The shortest turn in our study (0.6sec long) took much longer than the longest C start (0.21sec) in Domenici and Blake’s (1997) review. Our turns also have much lower maximum angular velocities, of around 600 deg/s, compared to 2000 deg/s or more in fast starts (Tytell and Lauder, 2008). Moreover, in fast starts, fish tend swing the head to one side (during the “C”) and back toward the other side as they kick out of the turn (Domenici and Blake, 1997), meaning that the head angular velocity has a positive and a negative peak. In our turns, the fish only swing their heads to one side, meaning that angular velocity only has a positive peak (Figure 3). During turning, bluegill in our study also reached a maximum snout linear velocity of 4.7 L/s, which is less than half of maximums that have been reported during bluegill C starts (Webb, 1978).

### But fast turns share some similarities with slow turns

Despite the discrete differences in timing and momentum, there are similarities between slow and fast turns. Whether they are performing a slow or fast turn, the turn begins with an anterior body movement, similar to what Gray (1933) saw in turning whiting. More recently, Thandiackal and Lauder (2020) found that the anterior and posterior body in zebrafish produce substantial positive work during turning, “planting” the tail and using it to drive the anterior body around through the turn.

Fast and slow turns are also quantitatively similar across a range of variables. First of all, regardless of whether bluegill turn slowly or quickly, they do not change how much they maximally bend their bodies, resulting in similar minimum moments of inertia (Figure 5). This may be a physical constraint based on the shape and thickness of the body.

Additionally, in both slow and fast turns, bluegill maximize their angular velocity around the same time that they minimize moment of inertia (Figure 7), similar to what Dabiri et al (Dabiri et al., 2020) found. Bluegill also consistently use a similar ratcheting behavior as tuna (Downs et al., 2023), suggesting that turning involves similar kinematics among different species.

### Pectoral fins and momentum may drive differences

There are both continuous and discrete differences in how bluegill use their pectoral fins during slow and fast turns. Pectoral fins play a greater role *continuously* during a slow turn, where fish tend to take more strokes, but a greater role *instantaneously* during a fast turn, where fish tend to take faster strokes that would produce higher torque (Figure 9). Additionally, many of these pectoral fin strokes are backing strokes. These occur when bluegill extend their inner fin far towards their heads during turning. These backing strokes are likely pushing water anteriorly to help the fish rotate. Drucker and Lauder (2001) quantified pectoral fin forces during turning bluegill and found that during turning pectoral fins act asymmetrically, and that both the turning side and weak side fins generate strong lateral and posterior forces respectively. Importantly, they studied bluegill turning slightly to avoid an obstacle (Drucker and Lauder, 2001) whereas we studied full 180 degree turns, suggesting that to turn more extremely, bluegill may also use their pectoral fins to generate backing forces. Finally, the backing behavior that we see has previously been observed in coelacanths who couple a backwards stroke of their inner pectoral fin with a forwards stroke of their outer pectoral fin during turning (Fricke and Hissmann, 1992).

Additionally, bluegill trade off linear and angular momentum differently in slow and fast turns. In both slow and fast turns, bluegill begin with little angular momentum and need to increase it to complete the turn (Figure 8 B, D). For fast turns, the angular momentum seems to come in part from their starting linear momentum, as linear momentum decreases at the same time that angular momentum increases (Figure 8 C, D). In contrast, during slow turns, bluegill do not start with as high linear momentum and subsequently increase both linear and angular momentum in order to turn around and accelerate in the opposite direction. This means they need to accelerate, gaining both linear and angular velocity.

### Future directions

We found that that fast turns are not sped up slow turns and are kinematically distinct, but there are some components that vary continuously. Therefore more work could be done to determine exactly when a slow turn becomes a fast turn. Is the transition gradual or discrete? Robotic and modelling studies could provide greater insight into this matter as they can directly control all features of how their “fish” turn and can test both the continuous and gradual paradigms.

While our work quantifies differences in slow and fast turns and highlights several mechanisms driving these differences, we are not able to parse apart the individual contributions of each fin. Therefore we advocate for detailed PIV during turning with a focus on what each fin is doing. In spontaneous routine turns, this is not really feasible, because fish may not turn consistently in the laser light sheet necessary for PIV. However, our setup means that fish turn in a controlled and repeatable way in a known position, which may make such flow measurements possible. Specifically, we show that bluegill use their pectoral and paired median fins extensively during turning. Therefore to better characterize turning and turning performance in fishes, we propose that future studies use PIV to focus on the pectoral and paired median fins during turning.

## ACKNOWLEDGEMENTS

We would like to acknowledge Dr. Barry Trimmer, Dr. Erik Dopman, Dr. Frank Fish, Dr. Yordano Jimenez, Dr. Mike Fath, and Dr. Ben Tidswell for their support and guidance on this project. We would also like to acknowledge the Tytell lab members for all of their help.

## COMPETING INTERESTS

The authors declare no competing interests.

## FUNDING

We received funding from NSF IOS 1652582 and ARO W911NF-17-1-0234.

## DATA AVAILABILITY

Code and data will be made available on Zenodo through Github upon publication.

## Notes

### Competing Interest Statement

The authors have declared no competing interest.

